# Population fragmentation leads to morpho-functional variation in British red squirrels (*Sciurus vulgaris*)

**DOI:** 10.1101/593319

**Authors:** Philip G. Cox, Philip J.R. Morris, Andrew C. Kitchener

**Affiliations:** Department of Archaeology and Hull York Medical School, University of York, Heslington, York, YO10 5DD, UK; School of Geosciences and Department of Integrative Biology, University of South Florida, 4202 E. Fowler Avenue, Tampa, FL 33620, USA; Department of Natural Sciences, National Museums Scotland, Chambers Street, Edinburgh, EH1 1JF, UK and Institute of Geography, School of Geosciences, University of Edinburgh, Drummond Street, Edinburgh EH9 3PX, UK

**Keywords:** Morphological evolution, masticatory biomechanics, mechanical advantage, geometric morphometrics, Sciuridae

## Abstract

It is well-known that population fragmentation and isolation can lead to rapid morphological and functional divergence, with the effect being particularly well-documented in rodents. Here, we investigated whether such a phenomenon could be identified in the Eurasian red squirrel (*Sciurus vulgaris*), which was once widespread across the majority of Great Britain, but suffered a severe population decline across the 20^th^ century, leaving a highly fragmented distribution. The aim was to test for morphological and biomechanical variation of the mandible between the remaining British red squirrel populations. Linear and geometric morphometric methods were used to analyse shape in a sample of over 250 squirrel mandibles from across the UK and Germany. Canonical variates analysis identified significant shape differences between most British red squirrel populations, but particularly between squirrels from Formby and those from other populations. Linear measurements showed that Formby red squirrels have a significantly lower mechanical advantage of the temporalis muscle, indicating that they are less efficient at gnawing. We suggest that this difference may be related to past supplemental feeding of Formby squirrels with peanuts, which are less mechanically resistant than food items that occur naturally in the diet of British red squirrels.

## Introduction

It is well-established that fragmentation of a population and the resulting isolation of the fragments can readily lead to morphological divergence. This effect has been noted in a wide variety of clades across the Animal Kingdom including butterflies (Hill et al. 1999), salmonid fishes (Snorrason, et al. 1994), lizards (Losos et al. 1997; Sumner et al. 1999), finches (Grant, 1999), and rodents (Renaud and Millien, 2001). Such divergence may represent the first step in allopatric speciation, the process by which a population splits into two reproductively isolated species as a result of an extrinsic barrier between them (Orr and Smith, 1998). The underlying mechanism leading to morphological divergence between isolated populations may be selective, i.e. an adaptive or plastic response to variations in local environmental pressures, or neutral, involving processes such as genetic drift or founder effect.

The rate of phentoypic change in isolated populations has been shown by many studies to be very rapid, with morphological variation following environmental change or introduction to a new habitat being detectable in just a few generations (Losos et al. 1997; Hale and Lurz, 2003; Kristjansson, 2005; Renaud et al. 2013; Holmes et al. 2016). In particular, island populations are known to undergo especially rapid morphological change, at least over short time periods (Millien, 2006), with the effect being particularly well-documented in rodents (Pergams and Ashley, 1999, 2001; Yom-Tov et al. 1999: Nargosen and Cardini, 2009). However, this effect is not restricted to islands and has also been shown to occur in mainland species subjected to habitat fragmentation (Schmidt and Jensen, 2003; Pergams & Lawler, 2009; Stumpp et al. 2018), indicating that many mammal species have the capacity to evolve at fast rates under changing environmental conditions (Millien, 2006). The most frequently reported rapid morphological changes are those of body size or mass (e.g. Schmidt and Jensen, 2003; Yom-Tov et al. 2008; Gardner et al. 2011). However, a number of studies have demonstrated that shape can also undergo change in a short period of time (e.g. Nargosen & Cardini, 2009; Franssen, 2011; Yazdi & Adriaens, 2011; Doudna and Danielson, 2015). The size and shape changes resulting from such rapid evolution on islands or in habitat fragments have been shown to have measurable functional consequences, particularly with regard to feeding biomechanics. For example, previous studies have demonstrated that morphological variation between closely-related insular species has resulted in different bite force capabilities in groups such as finches (Herrel et al. 2005), lizards (Herrel et al. 2008) and shrews (Cornette et al. 2012).

An ideal case study for studying the impact of population fragmentation and isolation is the British population of Eurasian red squirrels (*Sciurus vulgaris*). Once widespread across the majority of Great Britain (Shorten, 1954; Lloyd, 1983), the red squirrel began to suffer a severe population decline from the 1920s onwards (Gurnell, 1987). This has been attributed to various factors, such as loss of woodland habitat, competition with the introduced Eastern grey squirrel (*Sciurus carolinensis*), and parapoxvirus disease carried by grey squirrels (Tompkins et al. 2002; LaRose et al. 2010). Whatever the underlying reason for the population decline, it has resulted in a highly reduced and fragmented distribution of British red squirrels (Gurnell and Pepper, 1993; Barratt et al. 1999). Currently, red squirrels are found in most parts of Scotland except for the Central Belt, and across the northernmost counties of England, i.e. Northumberland and Cumbria (Gurnell et al. 2014). There are also isolated populations in County Durham, the Yorkshire Dales, the National Trust reserve at Formby on the Lancashire coast, and in some coniferous forests in mid Wales (Cartmel, 1997; Shuttleworth, 2000; Hobbs, 2005; Harris & Yalden, 2008). A population of red squirrels also existed in the lowland pine forest at Thetford in East Anglia until at least the early 21^st^ century (Gurnell et al. 2002: Rushton et al. 2002), but now appears to be extinct (Mathews et al. 2018). Beyond mainland Great Britain, red squirrels are found on Anglesey, the Isle of Wight and five islands in Poole Harbour, as well as on Jersey in the Channel Islands (Harris and Yalden, 2008; Simpson et al. 2010; Shuttleworth, 2010).

It should be noted that the remaining red squirrel populations in Britain are not all simply the relicts of a native subspecies distinct from red squirrels from mainland Europe. There have been numerous introductions of red squirrels from continental Europe over the last 150 years (Hale et al. 2004). In particular, the Jersey population was introduced from Europe, probably France, and southern England in the 1880s (Magris and Gurnell, 2001), and the Formby population was introduced from Europe, possibly Scandinavia, in the early to middle 20^th^ century (Lowe and Gardiner, 1983; Gurnell and Pepper, 1993). The red squirrels in the western half of northern England appear to be native, but in the east there have been some introductions from continental Europe (Hale and Lurz, 2003). Finally, the Scottish population seems to be mainly derived from re-introductions from England, but there have also been some introductions of individuals from Scandinavia (Harvie-Brown, 1880-1881).

The aim of this study is to determine whether the remaining isolated populations of British red squirrels have diverged in morphology from one another, and to assess the functional impact of any morphological differences. Specifically, it is hypothesised that differences in mandibular shape will be detectable between populations. This is predicted based on the phenomenon of rapid morphological change being well-known in rodents (Stumpp et al. 2018), and will be tested using geometric morphometrics (GMM; O’Higgins, 2000). It is also hypothesised that morphological variation in mandibular shape between squirrel populations will have a measurable impact on feeding biomechanics, as changes to the position of masticatory muscle insertions relative to the skull will impact the ability of those muscles to produce bite force. This second hypothesis will be tested using linear biomechanical measures.

## Methods

### Sample

The sample comprised 263 skeletonised red squirrel specimens from National Museums Scotland (Edinburgh, UK), most of which were collected between 1994 and 2006. Only specimens with associated location data were selected for this analysis. The sample included individuals from most areas of Great Britain and its offshore islands where red squirrels have been present over the last three decades, plus a number of specimens from Germany. The specimens chosen for analysis were grouped into the following geographical regions: North Scotland (Scotland north of the Central Belt); South Scotland (Scotland south of the Central Belt); North England (Northumberland and Cumbria); Formby; Thetford; Isle of Wight; Jersey; and Germany. Unfortunately, insufficient specimens from Wales were available for inclusion in the analysis. The number of specimens from each region is given in Table 1. From each specimen, one hemi-mandible was selected for analysis. Where both hemi-mandibles were present and undamaged, the right was used in preference to the left.

**Table 1.**
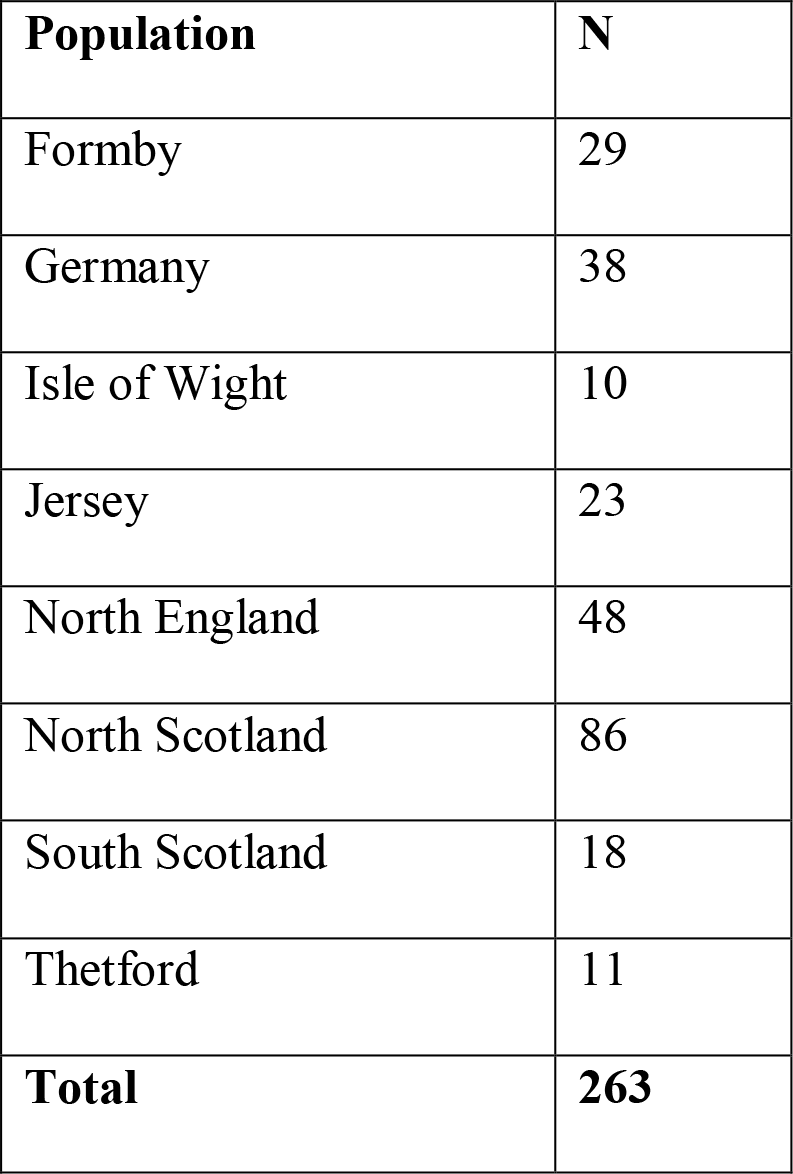
Number of specimens from each population in the analysis.

| Population | N |
| --- | --- |
| Formby | 29 |
| Germany | 38 |
| Isle of Wight | 10 |
| Jersey | 23 |
| North England | 48 |
| North Scotland | 86 |
| South Scotland | 18 |
| Thetford | 11 |
| **Total** | **263** |

### Morphometrics

Hemi-mandibles were laid flat with the external lateral surface facing upwards on paper marked with 0.5 mm squares (for scaling purposes). The specimens were photographed with a Panasonic Lumix DMC-TZ60 camera, secured on a tripod at a constant distance from the bench. A set of 12 two-dimensional landmarks, based on previous studies of rodent mandibles (e.g. Zelditch et al. 2008, 2015; Casanovas-Vilar and van Dam, 2013), was recorded from each photograph using the tpsDig2 software (Rohlf, 2018). The landmarks are illustrated and described in Figure 1A. Landmark co-ordinates from all 263 specimens were aligned via generalised Procrustes superimposition, and then subjected to a principal components analysis (PCA). Differences between populations were assessed using canonical variates analysis (CVA). Mahalanobis distances were calculated and the statistical significance of differences between groups was assessed with a permutation test of 10,000 repeats. All GMM analyses were carried out in MorphoJ (Klingenberg, 2011).

**Figure 1.**
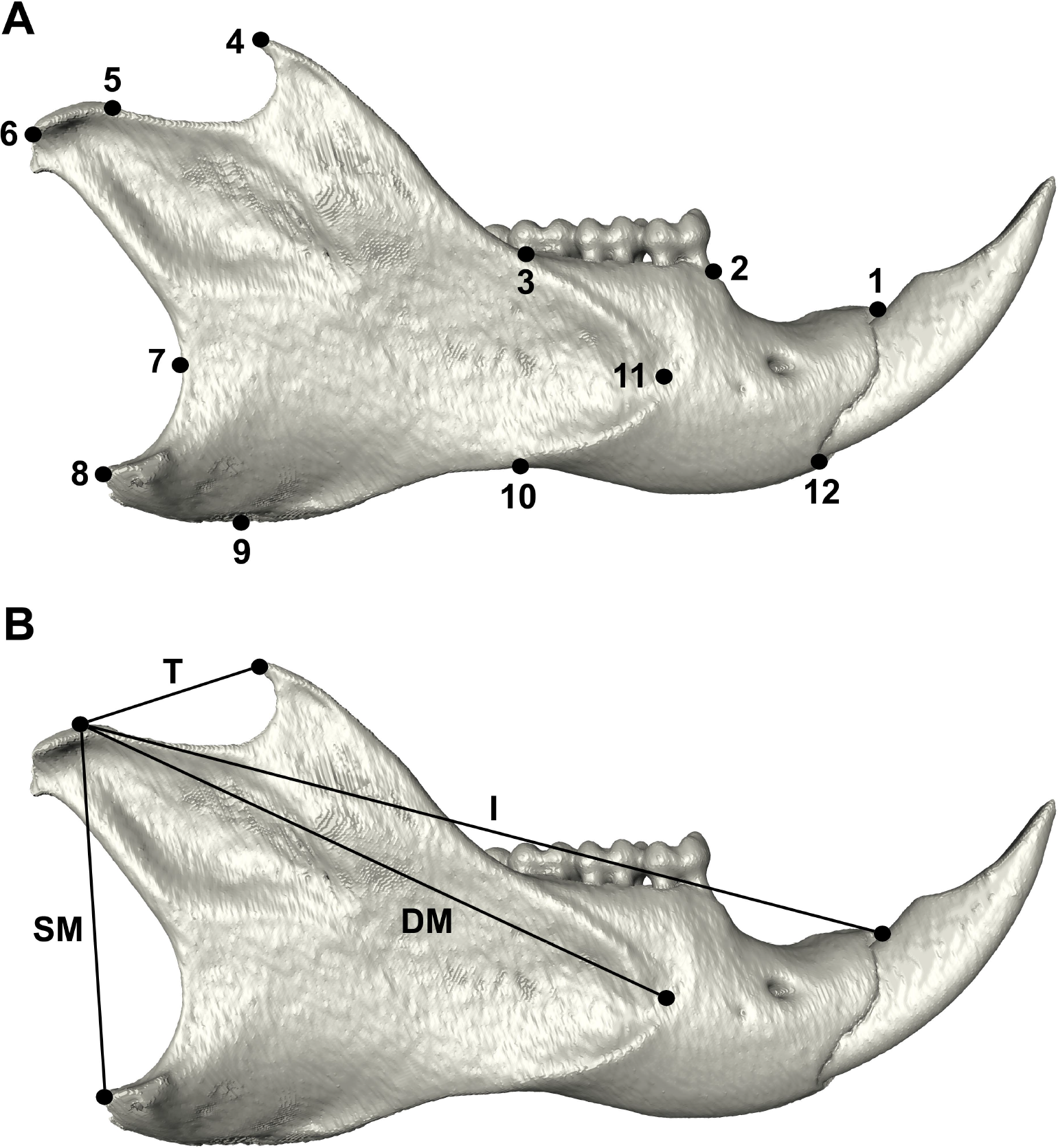
Right hemi-mandible of *Sciurus vulgaris* in lateral view showing (A) landmarks used in GMM and (B) in- and out-levers used in biomechanical analysis. Landmarks: 1, dorsalmost point on incisor alveolar margin; 2, anterior margin of premolar alveolus; 3, base of coronoid process where it crosses molar alveolar margin; 4, tip of coronoid process; 5, anteriormost point of condyle articular surface; 6, posteriormost point of condyle articular surface; 7, anteriormost point on curve between condyle and angular process; 8, posterior tip of angular process; 9, ventralmost point on angular process; 10, dorsalmost point on ventral border of ramus; 11, anteriormost point on masseteric ridge; 12, ventralmost point on incisor alveolar margin. Levers: DM, deep masseter in-lever; I, incisor out-lever; SM, superficial masseter in-lever; T, temporalis in-lever.

### Biomechanics

To elucidate the functional significance of any morphological variation between red squirrel populations, the mechanical advantage (MA) of three of the major jaw-closing muscles – temporalis, superficial masseter, and deep masseter – was estimated from linear measurements of the jaw (following Casanovas-Vilar and van Dam, 2013; Gomes Rodrigues et al. 2016). MA was calculated as the ratio of the muscle in-lever to the biting out-lever. To measure the lever lengths, an extra landmark was recorded at the dorsalmost point of the condyle articular surface (not included in the GMM analysis to avoid over-representing the condyle). The temporalis in-lever was measured from the condyle to the tip of the coronoid, the superficial masseter in-lever was measured from the condyle to the posterior tip of the angular process, and the deep masseter in-lever was measured from the condyle to the anteriormost point on the masseteric fossa margin. All muscle in-levers were compared to the out-lever representing incisor biting. As many of the specimens had missing or dislocated incisors, the out-lever was measured from the condyle to the dorsal margin of the incisor alveolus (as in Gomes Rodrigues et al. 2016). In and out-levers are illustrated in Figure 1B.

Muscle MAs were compared between red squirrel populations using ANOVA and post-hoc Tukey tests. The potential influence of size on MA was tested for using log centroid size calculated during the GMM analyses. All linear statistics were undertaken in PAST 3.06 (Hammer et al. 2001).

## Results

### Shape analysis

The distribution of individuals across the first two principal components, representing 24.9% and 18.7% of total variance respectively, is shown in Figure 2A. Little, if any, separation between the red squirrel populations is seen on these axes. PC1 represents changes to the morphology of the ventral margin of the mandible with mandibles at the positive end of the axis showing a more postero-ventrally deflected angular process. PC2 represents change in the body of the mandible from relatively shallow (negative) to deep (positive).

**Figure 2.**
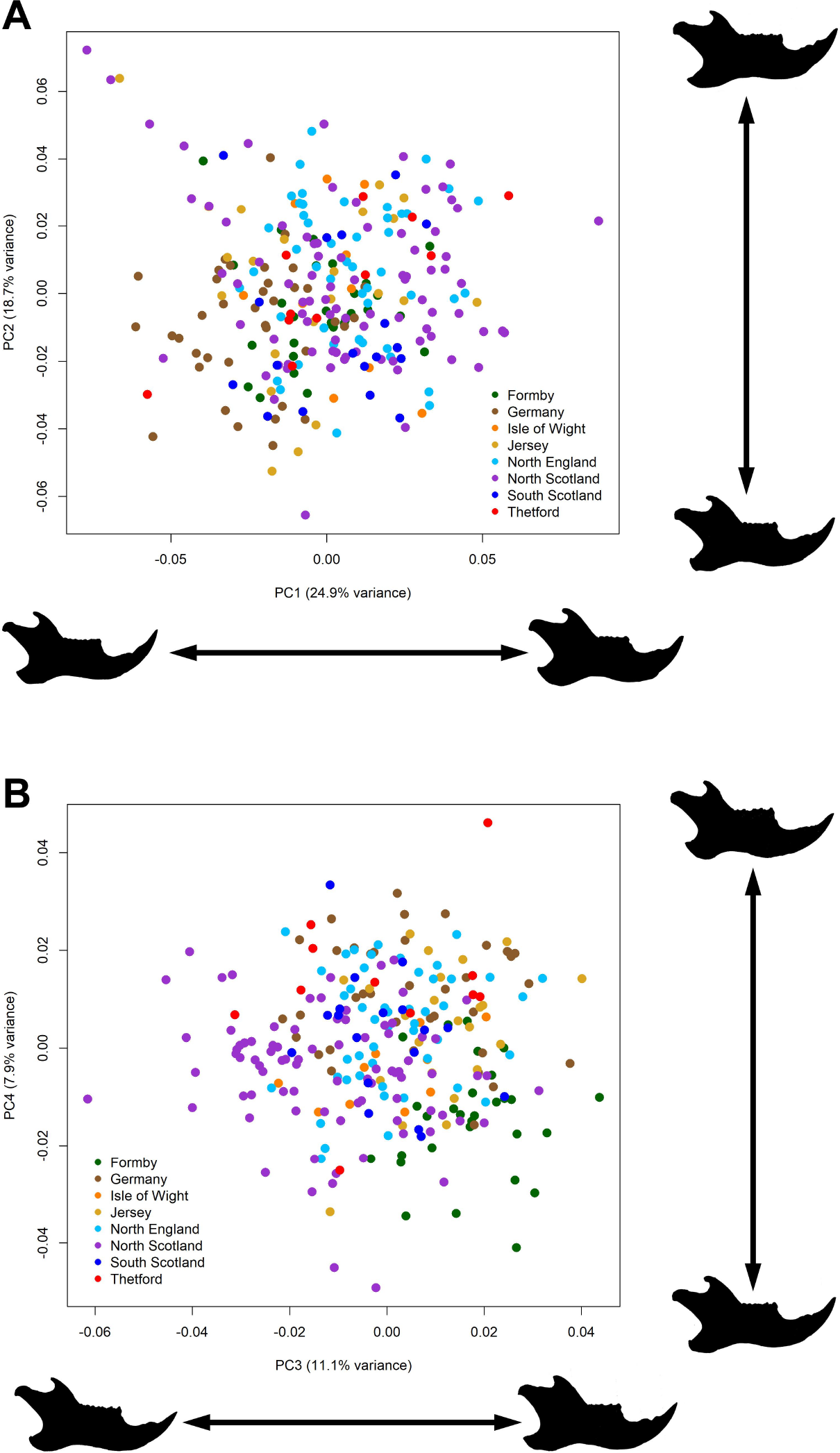
Principal components analysis of red squirrel mandibular shape. (A) PC1 versus PC2; (B) PC3 versus PC4.

Greater separation of squirrel populations is seen on the third and fourth PCs (11.1% and 7.9% total variance respectively), particularly with regard to the Formby population which is located in the positive PC3 and negative PC4 quadrants of Figure 2B. Mandibles at the negative end of PC3 have an anteriorly positioned coronoid and a relatively short angular process whereas mandibles with positive PC3 scores (such as the Formby squirrels) have a coronoid process that is more posteriorly positioned on the mandible and a more posteriorly extended angular process. Shape change along PC4 is more subtle, but again relates to the position of the coronoid process relative to the molar tooth row and condyle.

A plot of the first two canonical variates, together representing over 75% of total variance, is shown in Figure 3. It demonstrates a clear separation between many of the red squirrel populations, with the populations from Formby and Germany being particularly distinct. Permutation tests of 10,000 rounds indicate highly significant (*P* < 0.001) Mahalanobis distances between all pairs of red squirrel populations except between Northern England and Southern Scotland (*P* < 0.01), and the two smallest samples in this analysis, Thetford and the Isle of Wight (not significant). Mahalanobis distances and associated *P* values are given in Table S1.

**Figure 3.**
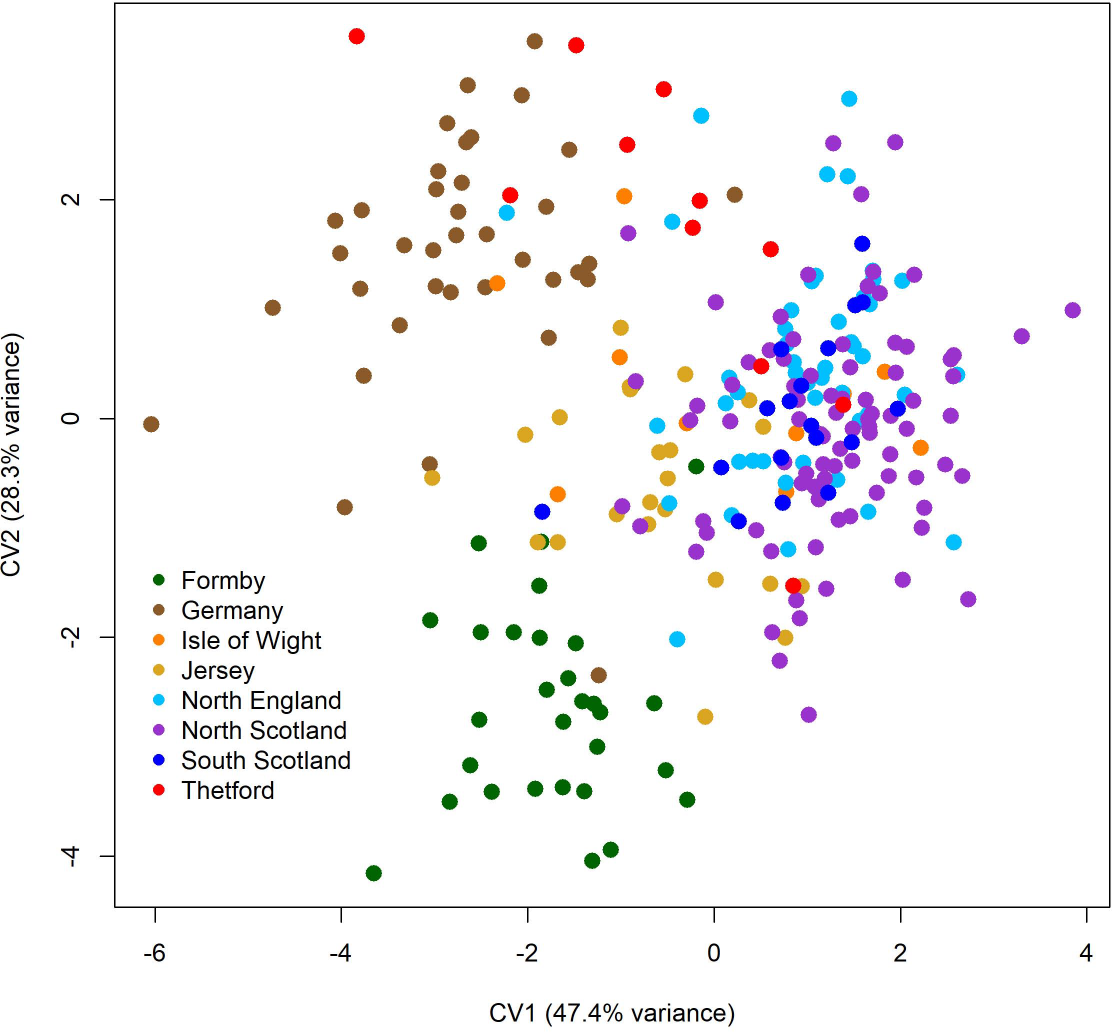
Canonical variates analysis of mandibular shape in red squirrel populations. First two CV axes displayed.

### Biomechanical analysis

The mean MA of each of the jaw-closing muscles in each of the red squirrel populations is given Table 2. Distributions within each population are shown in the boxplots in Figure 4. ANOVA tests revealed highly significant differences in MA between populations for the temporalis (*F* = 24.95, *P* < 0.001), superficial masseter (*F* = 3.09, *P* < 0.01), and deep masseter (*F* = 3.45, *P* < 0.01). It can be seen that the Formby squirrels have a much lower mean temporalis MA (0.181) than all other populations (0.205-0.225). Post-hoc pairwise Tukey tests confirmed significant differences between the Formby population and all other populations (Table S2). Significant differences were also revealed between the Jersey population and the Northern England, and both Scottish populations, and between the sample from Germany and the South Scotland and Northern England squirrels (Table S2). The mean MA for the superficial masseter ranged between 0.390 and 0.408 across different populations. No significant differences were found between pairs of populations except between those from Jersey and North Scotland (Table S3). The mean MA for the deep masseter ranged between 0.625 and 0.656. Significant differences were found between the German squirrels and the populations from Scotland and Northern England (Table S4).

**Table 2.** Mean MA ± standard deviation of masticatory muscles for red squirrel populations.

| Population | Deep masseter | Superficial masseter | Temporalis |
| --- | --- | --- | --- |
| Formby | 0.642 ± 0.018 | 0.402 ± 0.015 | 0.181 ± 0.015 |
| Germany | 0.625 ± 0.025 | 0.400 ± 0.021 | 0.210 ± 0.021 |
| Isle of Wight | 0.648 ± 0.024 | 0.390 ± 0.021 | 0.209 ± 0.011 |
| Jersey | 0.640 ± 0.032 | 0.408 ± 0.022 | 0.205 ± 0.014 |
| North England | 0.647 ± 0.025 | 0.400 ± 0.016 | 0.225 ± 0.015 |
| North Scotland | 0.647 ± 0.031 | 0.390 ± 0.024 | 0.217 ± 0.014 |
| South Scotland | 0.656 ± 0.028 | 0.398 ± 0.020 | 0.224 ± 0.016 |
| Thetford | 0.643 ± 0.021 | 0.402 ± 0.023 | 0.223 ± 0.016 |

**Figure 4.**
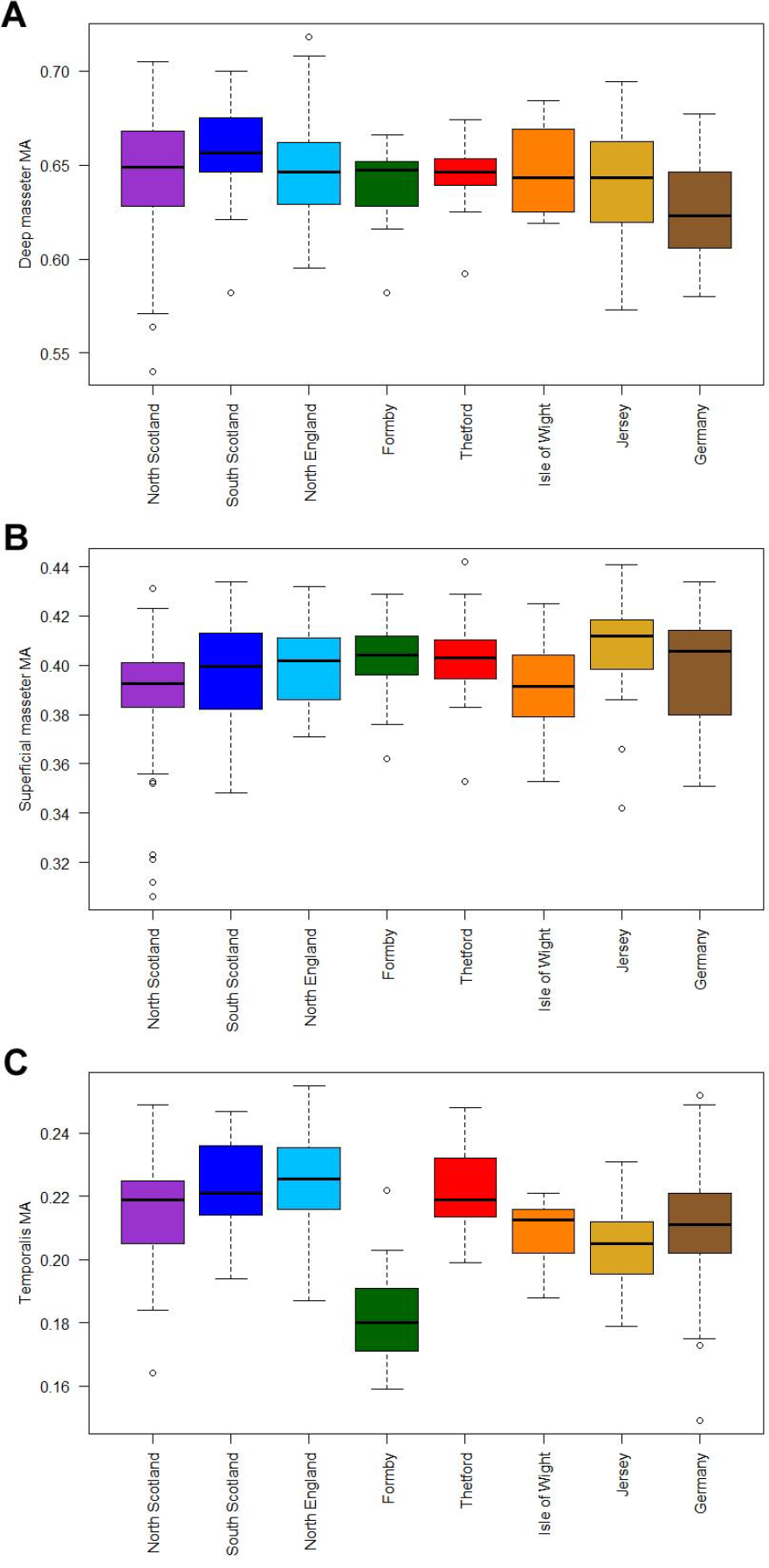
Boxplots of masticatory muscle MA in red squirrel populations. (A) deep masseter; (B) superficial masseter; (C) temporalis. Black line represents median, box represents interquartile range, open circles represent data points lying further than 1.5 times the interquartile range from the box.

An ANOVA revealed significant differences (*F* = 11.00, *P* < 0.001) in log centroid size of the mandible between squirrel populations. Post-hoc Tukey tests (Table S5) showed that the German squirrel mandibles are significantly larger than those of all other populations except Formby, and also that the Formby mandibles are significantly larger than those of all other populations except Germany and Jersey. Superficial masseter MA was found to be significantly positively correlated with size (*R* = 0.404, *P* < 0.001), as was deep masseter MA, although less strongly (*R* = 0.152, *P* < 0.05). The temporalis MA showed no significant correlation with mandibular centroid size (*R* = −0.109).

## Discussion

The decline of red squirrels in the UK and their replacement by invasive grey squirrels is a well-documented phenomenon (Lloyd, 1983; Gurnell, 1987). The current population of red squirrels in Great Britain is highly fragmented and mostly confined to northern areas and offshore islands (Gurnell and Pepper, 1993; Barratt et al. 1999; Mathews et al. 2018). The results of this study show that these isolated British red squirrel populations exhibit significant differences in mandibular morphology and mandibular function from one another, and from red squirrels from continental Europe. Notably, the red squirrel population from Formby, on the Lancashire coast, is shown to be particularly distinct from all other British populations, both in shape and biomechanical capabilities.

### Mandibular shape

The first hypothesis of this study, that differences in mandibular morphology between populations of red squirrels would be detectable with GMM, is supported clearly by the canonical variates analysis. CVA discriminates almost all populations from one another based on Mahalanobis distances (Table S1), with the separation of the Formby, Jersey and Germany populations being particularly evident on the plot of the first two canonical variates (Figure 3). This is perhaps not surprising as these three populations are each completely isolated from other red squirrel populations, whereas some of the other groupings defined in this analysis may not be completely separated from each other, e.g. North England, South Scotland and North Scotland. Furthermore, both the Formby and Jersey populations derive from continental Europe populations (Gurnell and Pepper, 1993; Magris and Gurnell, 2001), which may explain why they group with the German squirrels on CV1. In contrast, the PCA does not obviously discriminate between populations on the first two components (Figure 2A), although some separation can be seen on PC3 and PC4 (Figure 2B). The distribution of individuals on the first two principal components may be an artefact of the landmark set. Most of the shape change on PCs 1 and 2 is seen on the ventral margin of the mandible, specifically related to movements of landmarks 9 and 10. These two landmarks both represent maxima of curves and were noted to be more difficult to place reliably than other landmarks in the set. Future work with a set of semi-landmarks along the ventral mandibular margin may resolve this issue. On the third and fourth components, it is the Formby population that seems the most distinct in mandibular shape, as in the CVA. Examination of the shape changes along these PCs indicates that the position of the coronoid process relative to the condyle is the major morphological difference.

### Biomechanical performance

The second hypothesis, which predicted that differences in biomechanical performance would be detected between red squirrel populations, was also supported by the results here. The MA of the temporalis muscle in Formby red squirrels was shown to be significantly lower (*P* < 0.0001) than that of all other squirrel populations (Table S2). This aligns with the results from the shape analysis which indicated that the Formby squirrel mandibles have a coronoid process that is positioned closer to the condyle and further from the teeth. In addition, the MA of the temporalis MA in the Germany and Jersey populations was also shown to be significantly lower than some of the other squirrel populations, although to a much lesser extent. Analyses of centroid size showed that mandibles of the Formby and German squirrels were significantly larger than those of most other populations, but no significant correlation was found between centroid size and temporalis MA. Therefore, the reduced temporalis MA in these two populations does not appear to be an effect of allometric shape change.

No significant differences were detected in the MA of the superficial masseter muscle between populations, except in the singular case of the Jersey population versus the North Scotland population (Jersey squirrels have a higher superficial masseter MA than North Scotland squirrels). Similarly, only three significant differences in deep masseter MA were found, all between the German squirrels and other populations (Northern England and the two Scottish groupings). Although a positive correlation was found between deep masseter MA and size, the German squirrel mandibles had a significantly lower deep masseter MA, despite being significantly larger in size than all other populations. Thus, the functional differences found in German squirrel mandibles do not appear to be the result of allometric size change that is common to the whole sample.

A number of caveats to the biomechanical analysis should be noted to ensure that the results here are interpreted with appropriate levels of caution. Firstly, the use of lever arms to determine mechanical advantage will only give approximate results. To calculate MA more accurately, one should use moment arms i.e. the perpendicular distance between the condyle and the vector running between the muscle origin and insertion. This was not done as it would have required the skull and mandible to have been articulated in the photographs and intact skulls were not available for many of the specimens. Secondly, the representation of a muscle insertion as a single point is clearly inaccurate as the temporalis, superficial masseter and deep masseter all have large attachment sites on the squirrel mandible. Furthermore, all three muscles are complex and are divided into sections which have different fibre orientations and lines of action (Cox and Jeffery, 2011, 2015). Lastly, the analysis was conducted in 2D which ignores any lateral component to the mandible. Despite these simplifications to the biomechanical analysis, it was felt that the results generated were still meaningful, as the simplifications were consistent across all specimens and because squirrel hemi-mandibles are sciurognathous and therefore largely planar (Hautier et al. 2015).

### Formby red squirrels

The mandibles sampled from the red squirrel population in Formby were particularly distinct from those of other red squirrels in most of the analyses presented here. In particular, the Formby squirrels have been shown to have a much lower temporalis MA than all other British red squirrel populations (Figure 4C). The temporalis muscle in mammals is associated with fast closing of the jaws and a powerful bite force at the anterior teeth (Maynard Smith and Savage, 1959). Thus, the Formby squirrels appear to be less efficient at incisor gnawing than other squirrels in this sample – a situation that could be related to a number of (not necessarily mutually exclusive) factors.

First, it is important to note that the red squirrels at Formby appear not to be a fragment of the native British population, but originate from a European (possibly Scandinavian) population introduced to Ainsdale, Lancashire in the early 1930s (Lowe and Gardiner 1983; Gurnell and Pepper, 1993). Thus it might be suspected that the morphological distinctiveness of Formby red squirrels is a result of their different genetic background. However, this is probably too simplistic a view. The history of red squirrels in Britain has included numerous translocations from Europe over the last 200 years, leading to a complex phylogenetic relationship between current populations and no clear phylogeographical pattern (Barratt et al. 1999; Hale et al. 2004). A sample of red squirrels from Germany was included in this analysis to assess similarities between European and British populations. CVA distinguished the German squirrels from all other populations, and showed almost no overlap between the samples from Germany and Formby. Furthermore, the red squirrels from Germany did not show the highly reduced temporalis MA that was seen in the Formby squirrels.

Secondly, the red squirrel population at Formby is also known to be highly inbred, owing to the small founder population. This could have negatively affected the biomechanical capabilities of the mandible, as increased developmental instability and fluctuating asymmetry in the craniodental region has been suggested by some researchers to be associated with inbreeding and homozygosity (Leamy et al. 2002; Schaefer et al. 2006). However, others have found no evidence for this relationship (Markow, 1995), and a recent study on mice found no impact of inbreeding depression on bite force (Ginot et al. 2018).

Finally, the Formby red squirrel population, unlike all other populations in this analysis, is managed. The Sefton coast is a Special Area of Conservation and the pinewoods are a National Trust reserve for red squirrels. For several decades, the diet of the squirrel population at Formby was supplemented with peanuts provided by the National Trust and by the public (Gurnell & Pepper, 1993; Rice-Oxley, 1993; Shuttleworth, 2000), although this practice is now much reduced (A. Brockbank, pers. comm.). Peanuts are much less mechanically resistant than most of the food items that squirrels would naturally encounter in the UK (beech nuts, hazelnuts, conifer scales and seeds; Mollar, 1983). Thus it is possible that the morphology of the mandible, and hence the efficiency of gnawing, has changed in response to this change in diet. This explanation is consistent with the results seen here, as the less mechanically demanding peanuts could lead to a reduction in the MA of the temporalis, which is important for gnawing, but would have less effect on the MA of the superficial or deep masseter muscles, as they are more closely related to molar chewing. The morpho-functional change seen in Formby red squirrels could either be an evolutionary response that has occurred over a number of generations (as seen in insular populations e.g. Herrel et al. 2008; Cornette et al. 2012), or a plastic response that occurs across the lifetime of each individual exposed to supplemental feeding (as seen in laboratory animals raised on different diets e.g. He and Kiliaridis, 2003; Enomoto et al. 2010; Anderson et al. 2014). To tease apart these two possibilities would require a larger, well-dated sample of Formby red squirrel specimens spanning a number of decades.

The results generated by this study have important implications for conservation efforts related to British red squirrels. Garden feeding of red squirrels is popular amongst members of the public but may have unsuspected impacts on skeletal morphology if the food provided is less mechanically demanding than the squirrels would eat in the wild. Many conservation strategies in the UK currently involve translocation of squirrel individuals from well-populated areas or captive-breeding facilities to bolster threatened populations or initiate new ones. If it is true that supplemental feeding at Formby has led to changes in mandibular morphology that reduce the efficiency of gnawing, then diet must be taken into consideration during such translocations. The results here suggest that red squirrels may not thrive if moved to a habitat with a more mechanically demanding food source, or if supplementary feeding is withdrawn suddenly. This is consistent with previous research demonstrating that translocated red squirrels tend to survive longer in release sites that have a similar habitat to the origin site (Kenward and Hodder, 1998), and that animals released in unfamiliar habitat will tend to disperse away from the release site (Morris et al. 1993; Bright and Morris, 1994).

The conclusion that supplemental feeding has led to morphological change and the consequences this could have for conservation strategy is tantalising, but at the moment still preliminary. Future studies are planned that combine morphological with genetic data, that include data from the skull as well as the mandible, a longer time series of squirrels and that take advantage of more sophisticated biomechanical modelling techniques, in order to better understand the drivers of morpho-functional change in fragmentary populations such as British red squirrels.

## Author contributions

PGC conceived the study, photographed the mandibles and wrote the first draft of the manuscript. PJRM landmarked the specimens. ACK provided access to the specimens. All authors contributed to the final manuscript and approved its contents.

## Acknowledgements

We are grateful to the following people for helpful discussions on the analysis and interpretation of our data: Sam Cobb, Paul O’Higgins, Rob Ogden, Jeff Schoenebeck, and Mel Tonkin. We thank Andrew Brockbank of the National Trust for information on current squirrel management practices at Formby. PGC thanks Graham and Tis Buckley for accommodation in Edinburgh during data collection visits.

## Supplementary Information

**Table S1.**
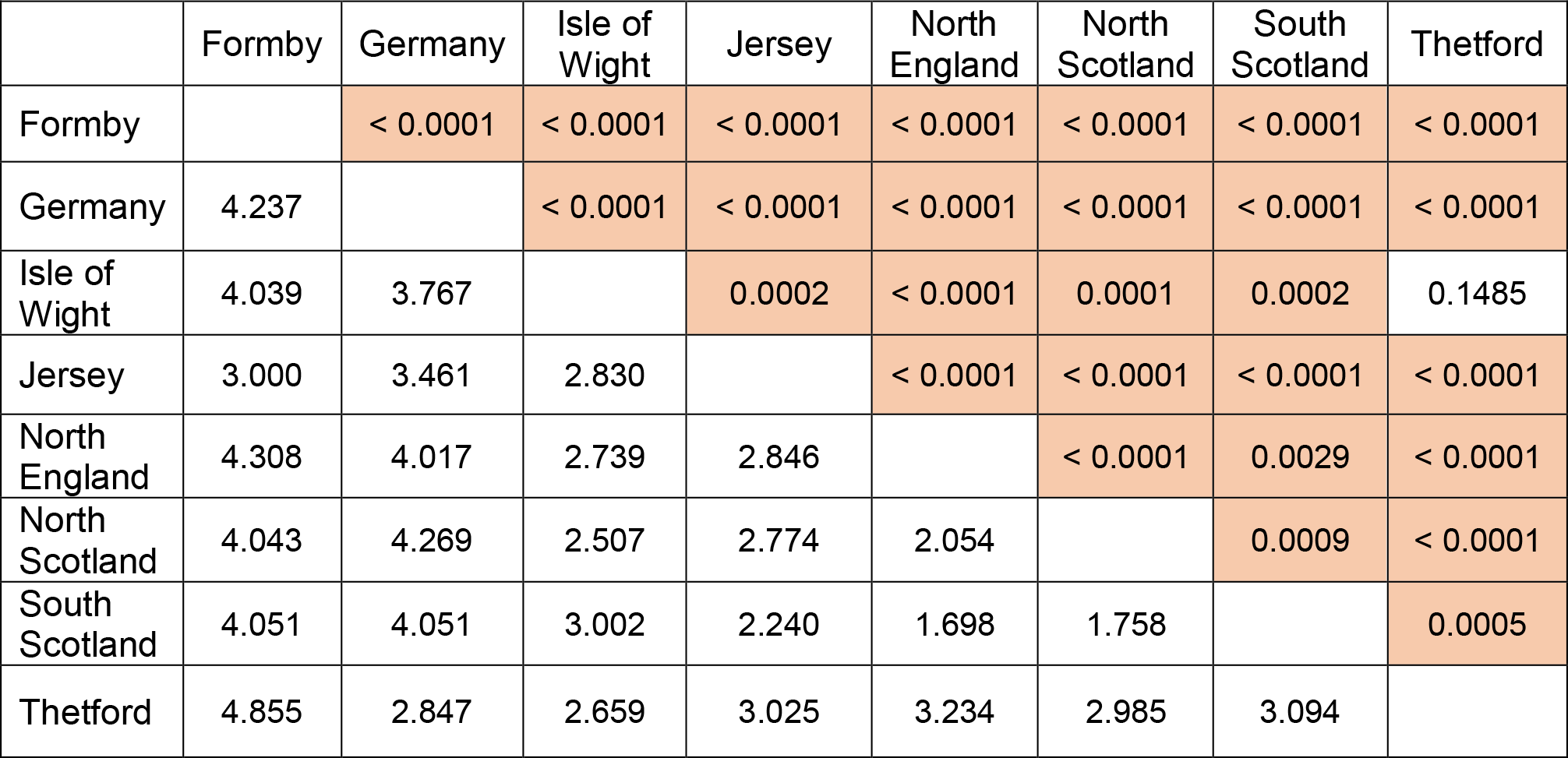
(A) Mahalanobis distances among squirrel populations (below diagonal) and *P* values from permutation tests (10000 permutation rounds) (above diagonal). Significant *P* values highlighted.

|  | Formby | Germany | Isle of Wight | Jersey | North England | North Scotland | South Scotland | Thetford |
| --- | --- | --- | --- | --- | --- | --- | --- | --- |
| Formby |  | < 0.0001 | < 0.0001 | < 0.0001 | < 0.0001 | < 0.0001 | < 0.0001 | < 0.0001 |
| Germany | 4.237 |  | < 0.0001 | < 0.0001 | < 0.0001 | < 0.0001 | < 0.0001 | < 0.0001 |
| Isle of Wight | 4.039 | 3.767 |  | 0.0002 | < 0.0001 | 0.0001 | 0.0002 | 0.1485 |
| Jersey | 3.000 | 3.461 | 2.830 |  | < 0.0001 | < 0.0001 | < 0.0001 | < 0.0001 |
| North England | 4.308 | 4.017 | 2.739 | 2.846 |  | < 0.0001 | 0.0029 | < 0.0001 |
| North Scotland | 4.043 | 4.269 | 2.507 | 2.774 | 2.054 |  | 0.0009 | < 0.0001 |
| South Scotland | 4.051 | 4.051 | 3.002 | 2.240 | 1.698 | 1.758 |  | 0.0005 |
| Thetford | 4.855 | 2.847 | 2.659 | 3.025 | 3.234 | 2.985 | 3.094 |  |

**Table S2.**
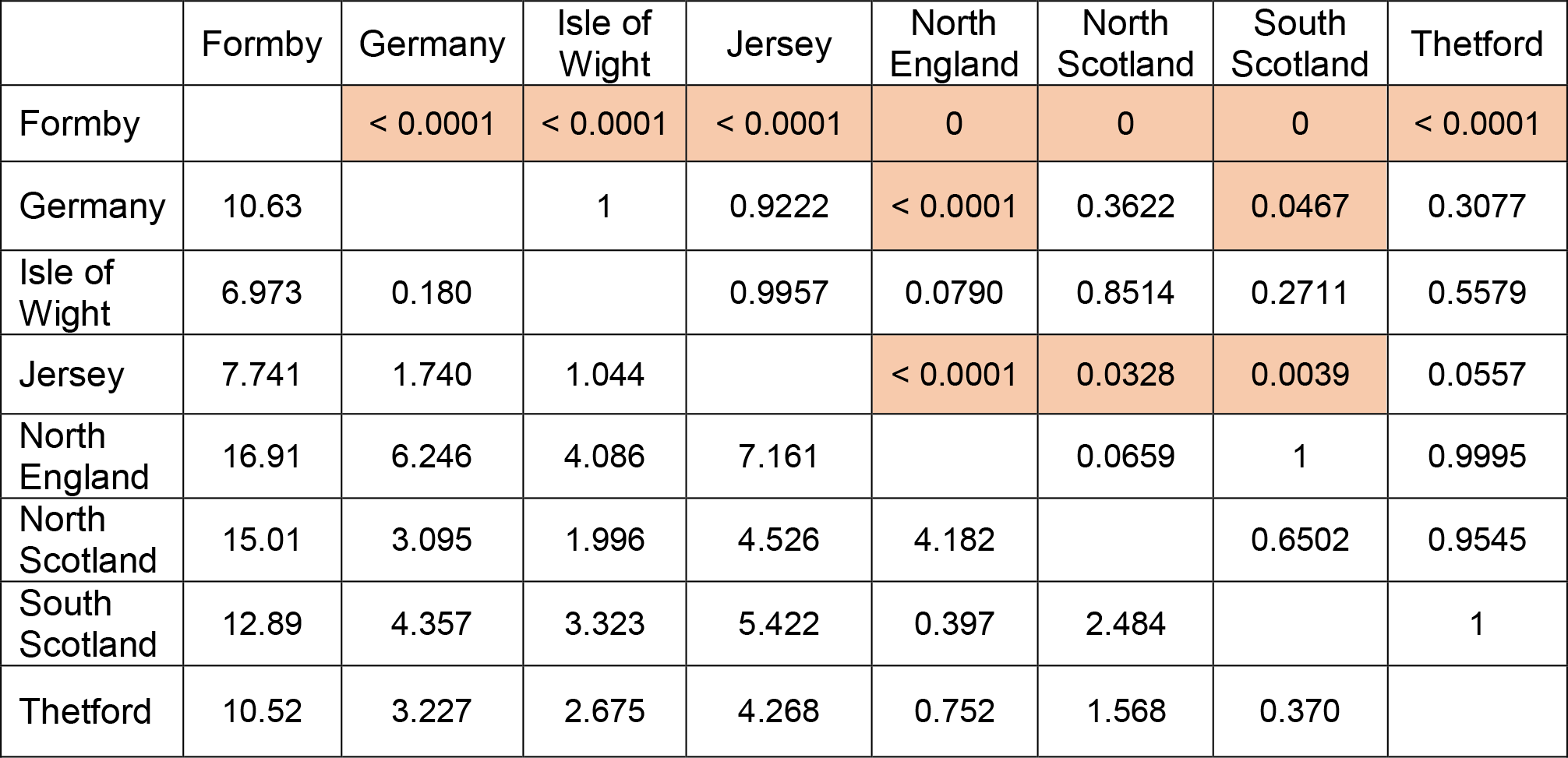
Post-hoc pairwise Tukey tests of temporalis MA between populations. Tukey’s Q statistic (below diagonal) and *P* values (above diagonal). Significant *P* values highlighted.

**Table S3.** Post-hoc pairwise Tukey tests of superficial masseter MA between populations. Tukey’s Q statistic (below diagonal) and *P* values (above diagonal). Significant *P* values highlighted.

|  | Formby | Germany | Isle of Wight | Jersey | North England | North Scotland | South Scotland | Thetford |
| --- | --- | --- | --- | --- | --- | --- | --- | --- |
| Formby |  | 1 | 0.7922 | 0.9725 | 1 | 0.1400 | 0.9986 | 1 |
| Germany | 0.353 |  | 0.8556 | 0.8976 | 1 | 0.1550 | 0.9999 | 1 |
| Isle of Wight | 2.160 | 1.983 |  | 0.3411 | 0.8595 | 1 | 0.9806 | 0.8811 |
| Jersey | 1.430 | 1.841 | 3.145 |  | 0.8634 | 0.0080 | 0.8146 | 0.9970 |
| North England | 0.432 | 0.071 | 1.971 | 1.959 |  | 0.1245 | 0.9999 | 1 |
| North Scotland | 3.762 | 3.700 | 0.047 | 5.142 | 3.832 |  | 0.8126 | 0.5739 |
| South Scotland | 0.873 | 0.611 | 1.344 | 2.101 | 0.570 | 2.106 |  | 0.9993 |
| Thetford | 0.108 | 0.366 | 1.900 | 0.985 | 0.420 | 2.642 | 0.784 |  |

**Table S4.** Post-hoc pairwise Tukey tests of deep masseter MA between populations. Tukey’s Q statistic (below diagonal) and *P* values (above diagonal). Significant *P* values highlighted.

|  | Formby | Germany | Isle of Wight | Jersey | North England | North Scotland | South Scotland | Thetford |
| --- | --- | --- | --- | --- | --- | --- | --- | --- |
| Formby |  | 0.2097 | 0.9974 | 1 | 0.9905 | 0.9891 | 0.6293 | 1 |
| Germany | 3.505 |  | 0.2352 | 0.4140 | 0.0068 | 0.0015 | 0.0020 | 0.5139 |
| Isle of Wight | 0.964 | 3.426 |  | 0.9928 | 1 | 1 | 0.9961 | 0.9999 |
| Jersey | 0.277 | 2.978 | 1.137 |  | 0.9748 | 0.9709 | 0.5673 | 1 |
| North England | 1.190 | 5.208 | 0.200 | 1.407 |  | 1 | 0.9300 | 0.9999 |
| North Scotland | 1.219 | 5.780 | 0.274 | 1.444 | 0.118 |  | 0.8763 | 0.9999 |
| South Scotland | 2.528 | 5.671 | 1.027 | 2.656 | 1.704 | 1.917 |  | 0.9165 |
| Thetford | 0.234 | 2.766 | 0.619 | 0.438 | 0.596 | 0.558 | 1.765 |  |

**Table S5.**
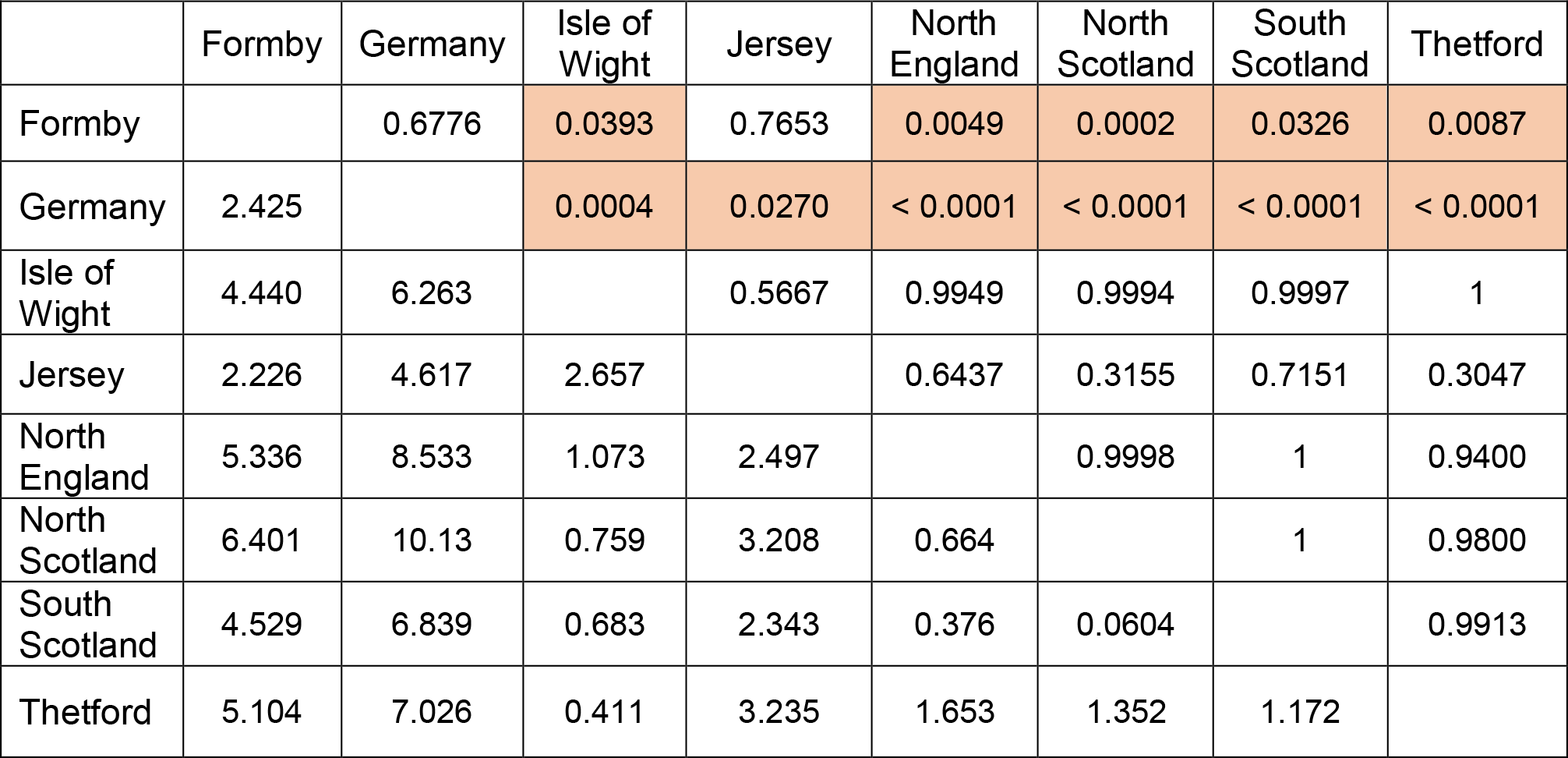
Post-hoc pairwise Tukey tests of log centroid size between populations. Tukey’s Q statistic (below diagonal) and *P* values (above diagonal). Significant *P* values highlighted.

|  | Formby | Germany | Isle of Wight | Jersey | North England | North Scotland | South Scotland | Thetford |
| --- | --- | --- | --- | --- | --- | --- | --- | --- |
| Formby |  | 0.6776 | 0.0393 | 0.7653 | 0.0049 | 0.0002 | 0.0326 | 0.0087 |
| Germany | 2.425 |  | 0.0004 | 0.0270 | < 0.0001 | < 0.0001 | < 0.0001 | < 0.0001 |
| Isle of Wight | 4.440 | 6.263 |  | 0.5667 | 0.9949 | 0.9994 | 0.9997 | 1 |
| Jersey | 2.226 | 4.617 | 2.657 |  | 0.6437 | 0.3155 | 0.7151 | 0.3047 |
| North England | 5.336 | 8.533 | 1.073 | 2.497 |  | 0.9998 | 1 | 0.9400 |
| North Scotland | 6.401 | 10.13 | 0.759 | 3.208 | 0.664 |  | 1 | 0.9800 |
| South Scotland | 4.529 | 6.839 | 0.683 | 2.343 | 0.376 | 0.0604 |  | 0.9913 |
| Thetford | 5.104 | 7.026 | 0.411 | 3.235 | 1.653 | 1.352 | 1.172 |  |

